# Antagonistic Roles of Tau and MAP6 in Regulating Neuronal Development

**DOI:** 10.1101/2024.02.21.581458

**Authors:** Xiaohuan Sun, Wenqian Yu, Peter W. Baas, Kazuhito Toyooka, Liang Qiang

## Abstract

Association of tau with microtubules causes them to be labile while association of MAP6 with microtubules causes them to be stable. As axons differentiate and grow long, tau and MAP6 segregate from one another on individual microtubules, resulting in the formation of stable and labile domains. The functional significance of the yin/yang relationship between tau and MAP6 remained speculative in those studies, with one idea being that such a relationship assists in balancing morphological stability with plasticity. Here, using primary rodent neuronal cultures, we show that depletion of tau has opposite effects compared to depletion of MAP6 on the rate of neuronal development, the efficiency of growth cone turning, and the number of processes and axonal branches. Opposite effects to those of tau depletion were also observed on the rate of neuronal migration, in an *in vivo* assay, when we depleted MAP6. When tau and MAP6 were depleted together in the cell culture assays, the morphological phenotypes negated one another. Tau and MAP6 are multifunctional proteins, but the present results suggest that the observed effects of their depletion on neuronal development are likely due to their opposite roles in regulating microtubule dynamics.

**Summary:** Tau and MAP6 play antagonistic roles in regulating multiple aspects of neuronal development, presumably via their antagonistic effects on microtubule dynamics.

## Introduction

Microtubules (MTs) serve as the scaffold of neuronal architecture (Kapitein and Hoogenraad 2015, Baas, Rao et al. 2016). In the axon, each MT consists of a stable domain toward its minus end and a labile domain toward its plus end. The former is characterized by reduced tubulin subunit exchange and greater resistance to depolymerization, whereas the latter is prone to rapid polymerization and depolymerization, reflecting a higher dynamicity (Baas and Black 1990, Baas, Rao et al. 2016). The properties of these domains derive from their association with various binding partners, including MT-associated proteins (MAPs), and are reflected in their levels of tubulin post-translationally modified by detyrosination and acetylation (Brown, Li et al. 1993, Sudo and Baas 2010, Janke and Magiera 2020). Through the interplay of different binding partners, the two MT domains can vary in their degree of stability/dynamicity so that the labile domain can be more or less labile, and the stable domain can be more or less stable (Baas, Rao et al. 2016, Qiang, Sun et al. 2018, Baas and Qiang 2019). Whether individual MTs have such distinct stable and labile domains elsewhere in the neuron remains unclear, but stable and labile MT fractions have been shown to exist throughout the life of the neuron and in different neuronal compartments (Baas, Rao et al. 2016).

Tau, an abundant MAP encoded by the MAPT gene on human chromosome 17, is well known for influencing the properties of MTs in the axon and, in turn, the morphology of the axon (Arendt, Stieler et al. 2016, Wang and Mandelkow 2016, Mueller, Combs et al. 2021). Long-standing dogma implicates tau in binding to axonal MTs to enhance their stability (Dehmelt and Halpain 2004, Kolarova, García-Sierra et al. 2012, Kadavath, Hofele et al. 2015), but this view was challenged by our recent studies on cultured rodent neurons using RNA interference (RNAi) to reduce tau levels. These studies showed that reducing tau levels did not result in MT destabilization in the axon but rather in the selective loss of the labile MT fraction, with the remainder of that fraction becoming more stable (Qiang, Sun et al. 2018). Additional work showed that heightened stabilization of the remainder of the labile fraction was due to increased binding of another MAP, namely MAP6, previously termed Stable-Tubule-Only-Polypeptide (STOP). MAP6, a protein with documented MT-stabilizing properties, is enriched on the stable domains of axonal MTs (Slaughter and Black 2003, Cuveillier, Boulan et al. 2021). Taken together, these findings suggest that tau enables the assembly of long labile domains of MTs in the axon and prevents them from being stabilized by MAP6, at least until a signaling pathways recalibrates the balance between these two MAPs to allow MT stabilization to occur.

The balance between stable and labile MT fractions has long been recognized as significant to diverse cellular processes, including axonal growth and branching, fast axonal transport, growth cone navigation, neuronal migration, and synaptic homeostasis (Baas, Rao et al. 2016). Investigating this with regard to tau and MAP6 is challenging because each of these two MAPs is multi-functional with complex interactions and responsibilities (Lefèvre, Savarin et al. 2013, Gory-Fauré, Windscheid et al. 2014, Tortosa, Adolfs et al. 2017, Cuveillier, Delaroche et al. 2020, Cuveillier, Boulan et al. 2021, Mueller, Combs et al. 2021, Bloom and Baas 2024). However, if depletion of each of them individually produces phenotypes that are the opposite to one another, such effects would likely be due to their opposite effects on MT dynamics. In this short report, we have sought to test this prediction in three different experimental systems.

## Results and Discussion

### Tau and MAP6 reductions have opposite effects on tyrosinated tubulin levels, neuronal development, and morphology

A variety of studies on neurons from our group have consistently documented a strong correlation between the content of tyrosinated tubulin in MTs and their dynamicity (Brown, Li et al. 1993, Baas, Rao et al. 2016). In growing axons, the levels of tyrosinated tubulin in the labile domain of the MTs is high while the stable domain shows little or no labeling for tyrosinated tubulin in studies using either immuno-electron or immunofluorescence microscopy (Baas and Black 1990, Brown, Li et al. 1993). In the distal region of the axon, the labile domains are especially labile, and correspondingly, their levels of tyrosinated tubulin are higher yet than in the labile domains along the axon shaft (Ahmad, Pienkowski et al. 1993). Thus, as a general principle, the levels of tyrosinated tubulin serve as a marker for how labile the MTs are in a particular MT fraction analyzed.

After confirming the effectiveness of tau or MAP6 knockdown via western blotting in hippocampal neurons after 4 days of siRNA interference (Figure 1A-B), we then examined tyrosinated tubulin levels and found a 31.88% decrease compared to control neurons when tau was knocked down. Conversely, a 30.01% increase was observed in MAP6-reduced neurons compared to controls (Figure 1C-D). These findings are consistent with our previous work showing that tau reduction leads to diminution of the labile fraction, whereas MAP6 reduction increases the labile fraction (Qiang, Sun et al. 2018). Moreover, the quantitatively opposite effects on tyrosinated tubulin levels are well suited to test our hypothesis about a potential yin/yang relationship between tau and MAP6, via their antagonistic effects on MT dynamics. It is worth noting that when tau is depleted, MAP6 levels rise, while conversely, depletion of MAP6 results in elevated tau levels (Figure 1A-B). This indicates that the levels of tau and MAP6 react to one other. Next, we used hippocampal neuron cultures to explore the impact of tau and MAP6 reductions on neuronal morphology during their development. Similar to the above, siRNA of tau and MAP6 were employed to reduce the expression of tau, MAP6, or both in combination. These neurons were cultured for 2 days after nucleofection with the corresponding siRNA and then replated and cultured for an additional 2 days. We examined neuronal morphology by categorizing neurons based on their developmental stages (Figure 1E), following the previously published classification (Dotti, Sullivan et al. 1988). Briefly, in stage I, neurons exhibit a lack of processes and maintain a rounded shape encircled by lamellipodia. Moving to stage II, neurons begin to develop a few minor processes around the cell body, and in stage II, one of these minor processes extends to become the axon. Our results show that 4.9% more tau knockdown neurons stay in Stage I, while 4.9% fewer neurons achieve Stage II and III compared to control neurons, indicating that the reduction of tau slightly slows neuronal development compared to hippocampal neurons treated with the control siRNA (Figure 1F). On the other hand, MAP6-reduced neurons display the opposite phenotype, with accelerated neuronal development (Figure 1F), showing that 5.2% more tau knockdown neurons achieve Stage II and III, while 5.2% fewer neurons stay in Stage I compared to control neurons. These findings indicate that tau reduction in cultured hippocampal neurons delays their stepwise development, whereas MAP6 reduction accelerates it. Strikingly, no significant difference was found in neuronal developmental stages between control siRNA-treated neurons and those in which both tau and MAP6 were simultaneously knocked down via siRNA (Figure 1F).

**Figure 1:**
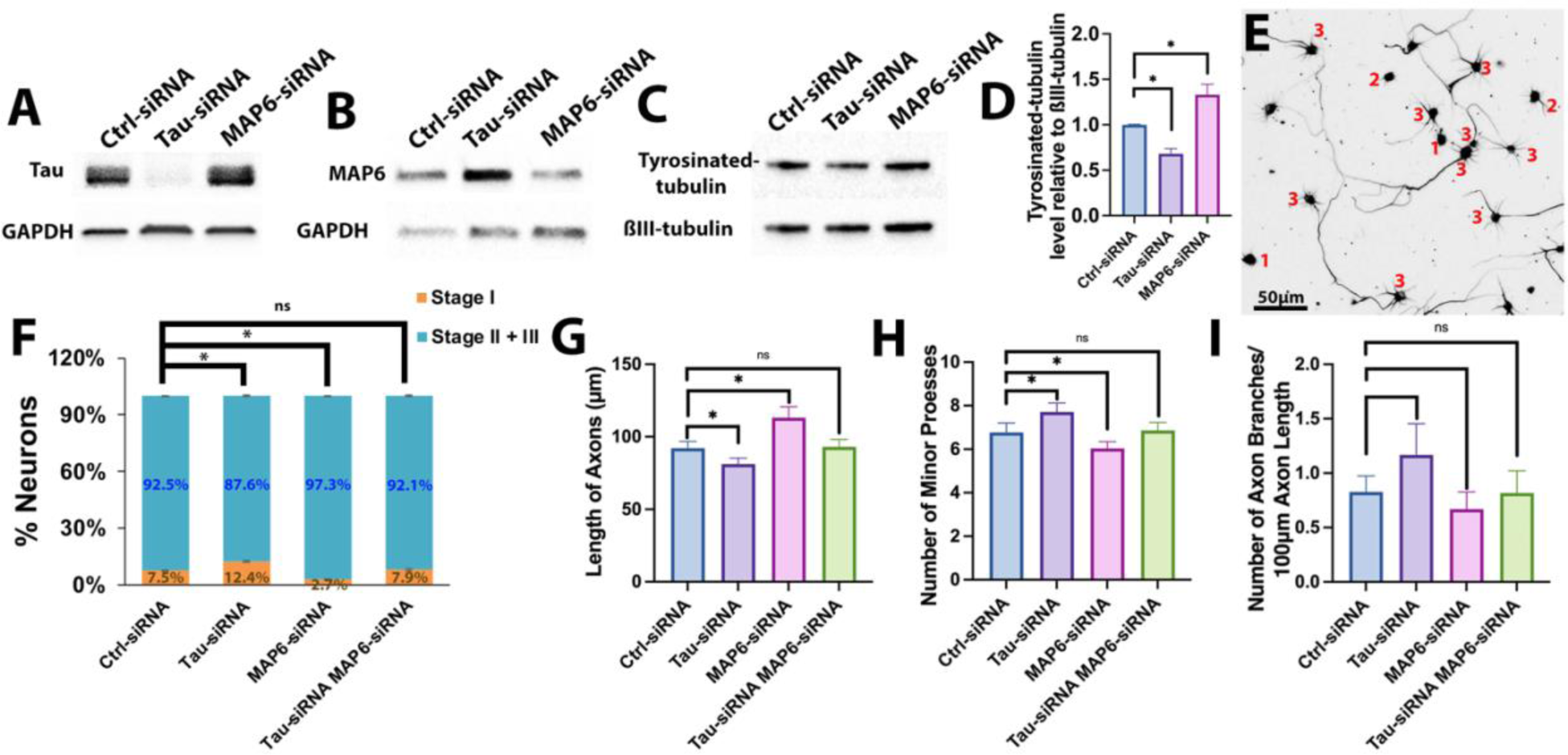
Opposite effects of depletion of tau or MAP6 on various aspects of MT dynamics and neuronal development in rat hippocampal neurons. (A-B) Representative western blots depicting tau levels, MAP6 levels, and corresponding GAPDH levels in control hippocampal neurons, as well as in neurons treated individually with tau siRNA and MAP6 siRNA. (C) Representative western blots depicting tyrosinated tubulin levels and ßIII-tubulin levels in control, tau siRNA, and MAP6 siRNA-interfered hippocampal neurons. (D) The bar graph depicts tyrosinated tubulin intensity relative to ßIII-tubulin intensity ± SEM for control siRNA (n=3), tau siRNA (n=3), and MAP6 siRNA (n=3). Hippocampal neurons treated with tau siRNA showed a 31.88% ± 5.70% lower tyrosinated tubulin intensity compared to control neurons (100% ± 0.64%) (one-way ANOVA \**p* < 0.05). In contrast, neurons treated with MAP6 siRNA exhibited a 33.01% ± 11.66% higher tyrosinated tubulin intensity compared to control neurons (100% ± 0.64%) (one-way ANOVA \**p* < 0.05). (E) Inverted fluorescence image displaying cultured hippocampal neurons stained with ßIII tubulin in stages I (1), II (2), and III (3), respectively. Scale bar = 50µm. (F) A stacked graph represents the percentage of neurons in Stage I and combined Stages II and III in control siRNA (n=135), tau siRNA (n=147), MAP6 siRNA (n=145), and neurons simultaneously interfered with both tau siRNA and MAP6 siRNA (n=102). When tau is knocked down, there is a significant increase of 4.90% ± 0.55% in the proportion of neurons remaining in Stage I compared to control neurons (7.46% ± 0.20%) (t-test \**p* < 0.05). Conversely, when MAP6 is knocked down, there is a significant decrease of 4.76 ± 0.05% in the proportion of neurons remaining in Stage I compared to control neurons (7.46% ± 0.20%) (t-test \**p* < 0.05). However, there is no significant difference (ns) observed between control neurons and the neurons that were interfered with both tau and MAP6 siRNA (7.92% ± 0.53%), simultaneously. (G-I) The bar graphs illustrate measurements for axon length, number of minor processes, and number of axon branches per 100µm of axon length in different experimental conditions: control neurons, neurons with tau siRNA interference, neurons with MAP6 siRNA interference, and neurons with both tau siRNA and MAP6 siRNA interference. (G) The bar graph displays axon length ± SEM for control siRNA (n=50), tau siRNA (n=59), MAP6 siRNA (n=47), and tau and MAP6 siRNA (n=51). Tau siRNA resulted in a 10.97 ± 4.12 µm shorter axon compared to the control axon (92.14 ± 4.73 µm) (t-test, \**p* < 0.05). Conversely, MAP6 siRNA exhibited a 21.01 ± 7.54 µm longer axon compared to control siRNA (92.14 ± 4.73 µm) (t-test, \**p* < 0.05). No significant difference (ns) was observed between the control neurons and those treated with both tau and MAP6 siRNA (93.03 ± 5.12 µm). (H) The bar graph quantifies the number of minor processes ± SEM for control siRNA (n=66), tau siRNA (n=65), MAP6 siRNA (n=60), and tau and MAP6 siRNA (n=64). Tau siRNA showed a 1.21 ± 0.47 higher number of minor processes compared to the control (6.95 ± 0.47) (t-test, \**p* < 0.05). On the other hand, MAP6 siRNA exhibited a 0.69 ± 0.40 lower number of minor processes compared to control siRNA (6.95 ± 0.47) (t-test, \**p* < 0.05). There was no notable difference (ns) observed between the control neurons and neurons were transfected simultaneously with tau and MAP6 siRNA (6.86 ± 0.37). (I) The bar graph quantifies the number of axon branches per 100µm of axon length ± SEM for control siRNA (n=49), tau siRNA (n=33), MAP6 siRNA (n=48), and tau and MAP6 siRNA (n=38). Tau siRNA showed a 0.31 ± 0.29 higher number of axon branches per 100µm of axon length compared to the control (0.85 ± 0.17) (t-test \**p* < 0.05). MAP6 siRNA exhibited a 0.19 ± 0.16 lower number of axon branches per 100µm of axon length compared to control siRNA (0.85 ± 0.17) (t-test \**p* < 0.05). There was no notable difference (ns) observed between the control neurons and neurons were transfected simultaneously with tau and MAP6 siRNA (0.81 ± 0.20).

Next, we evaluated the length of the axon, the number of minor processes, and the number of axonal branches per 100 µm in stage 3 neurons. Compared to the control, tau-reduced neurons developed shorter axons by 10.97 µm with more minor processes by 1.21 per soma (Figure 1G-H). In contrast, MAP6-reduced neurons extended longer axons by 21.01 µm with fewer minor processes by 0.69 per soma, compared to control neurons (Figure 1G-H). Indeed, tau-reduced neurons had greater numbers of axonal branches by 0.31 per 100 µm in axonal length, compared to control neurons; by contrast, MAP6-reduced neurons had fewer axonal branches by 0.19 per 100 µm, compared to control (Figure 1I). When tau and MAP6 were simultaneously knocked down via siRNA, no differences in axon lengths, the numbers of minor processes, or the numbers of axonal branches when compared to the control siRNA treated neurons (Figure 1G-I).

### Tau and MAP6 reductions have opposite effects on axonal navigation

To investigate the influence of tau and MAP6 on growth cone turning, we carried out experiments using a previously established laminin/poly-D-lysine border assay. For these studies, we used SCG neurons because they were previously used with this assay (Nadar, Ketschek et al. 2008, Liu, Nadar et al. 2010), and because we wished to test our hypothesis about tau and MAP6 in different kinds of neurons as well as challenged in different ways. In this assay, these neurons were cultured for 2 days after nucleofection with the corresponding siRNA on a platform coated with poly-D-lysine, with Laminin added to the medium. They were then replated and cultured for an additional 2 days on the same platform, with one-half coated with poly-D-lysine and the other half coated with a laminin matrix layered on top of the poly-D-lysine. The laminin-coated section is the preferred growth environment, whereas the other half lacks laminin, creating an environment much less preferred. As their axons extend and reach the laminin/poly-D-lysine border, owing to their inclination to remain on the laminin side, most growth cones guide the axons to turn and persist on the laminin-coated region. This process requires labile domains of MTs extending into the distal segment of the axon and requires them to be even more labile than the labile domains in the axon shaft (Buck and Zheng 2002, Suter, Schaefer et al. 2004). Strikingly, when tau levels were reduced by siRNA, we observed a 50.03% decrease in the amount of SCG axons turning towards the laminin border, compared to control neurons. By contrast, when MAP6 was reduced, we observed a 23.49% increase in the amount of SCG axons turning towards the laminin border, compared to control neurons. Notably, the simultaneous knockdown of tau and MAP6 through siRNA were indistinguishable from control neurons with regard to growth cone turning (Figure 2A-E).

**Figure 2:**
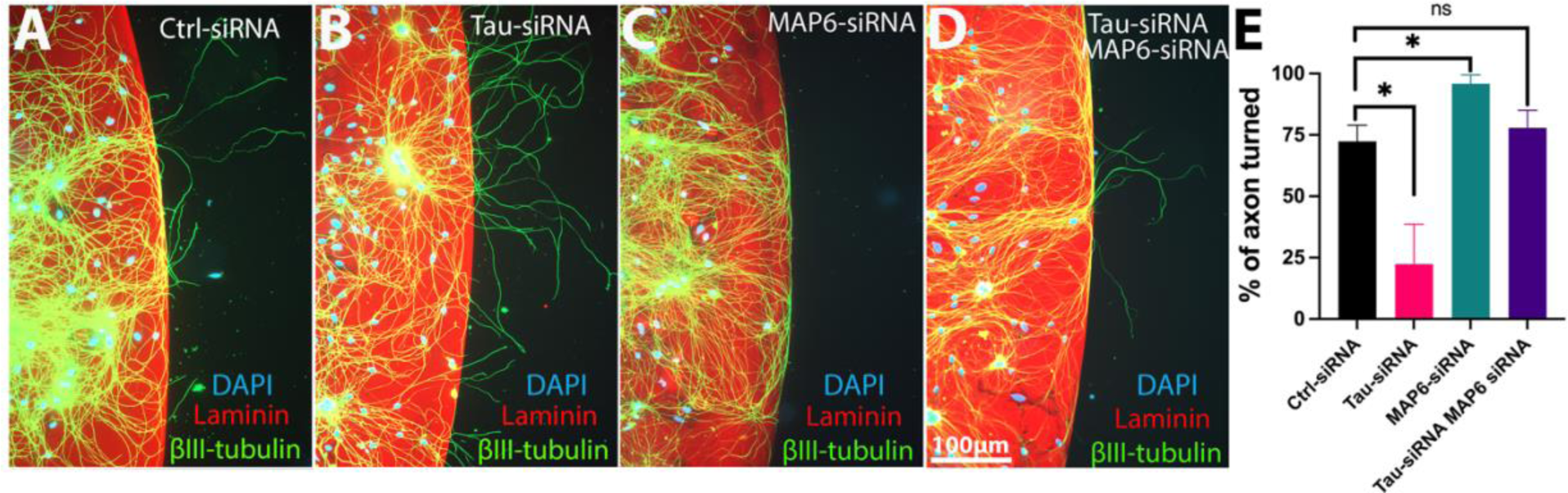
Opposite effects of depletion of tau or MAP6 on growth cone turning of cultured rat SCG neurons. (A-D) Fluorescence images of SCG neurons transfected with control siRNA, tau siRNA, MAP6 siRNA, or both tau and MAP6 siRNA simultaneously, with DAPI staining in blue, laminin in red, and ßIII-tubulin in green. Scale bar = 100 microns. (E) A bar graph illustrates the quantification of the percentage of axons turning at the Laminin/Poly-D-Lysine border ± SEM for control siRNA (n=95), tau siRNA (n=97), MAP6 siRNA (n=109), and both tau and MAP6 siRNA (n=89), where ‘n’ represents the number of axons near the Laminin/Poly-D-Lysine border, including both crossing and turning at the border. Tau siRNA resulted in a 50.03% ± 9.39% lower percentage of axons turning around the Laminin/Poly-D-Lysine border compared to the control (72.37% ± 3.78%) (one-way ANOVA, \**p* < 0.05). Conversely, MAP6 siRNA showed a 23.49% ± 2.08% higher percentage of axons turning around the Laminin/Poly-D-Lysine border compared to the control (72.37% ± 3.78%) (one-way ANOVA, \**p* < 0.05). There was no notable difference (ns) observed between the control neurons (72.37% ± 3.78%) and those that were transfected simultaneously with tau and MAP6 siRNA (77.91% ± 4.12%).

### Tau or MAP6 depletion have inverse effects on the rate neuronal migration

Cortical neurons are generated in the ventricular zone (VZ) and migrate to their destinations in an inside-out fashion (Stiles and Jernigan 2010). The migration process involves leading process extension, nucleus translocation, and tail process retraction (Lambert de Rouvroit and Goffinet 2001). This is crucial for neuronal development as it ensures precise cell positioning, maintains spatial organization, and facilitates proper synaptic connections (Marín, Valiente et al. 2010). Previous studies demonstrated that tau reduction through shRNA results in tardiness in migrating neurons reaching their destinations (Sapir, Frotscher et al. 2012). To investigate whether MAP6 has an inverse effect on neuronal migration, we conducted a similar experiment by injecting specific MAP6 shRNA into mouse embryos at E15.5 (Figure 3A). At P0, the brains were collected for fixation and analysis. The effectiveness of MAP6 knockdown was verified (Figure 3B). Then we conducted an analysis of neuronal migration speed by dividing the cortical region into 5 bins of equal size. Subsequently, we examined the percentage of neurons within each bin. A higher concentration of neurons in bin #5 denotes faster migration. Conversely, a larger population of neurons remaining in bin #1 suggests slower migration. Our results show that successfully transfected neurons whose MAP6 levels are reduced were labeled with Venus. 10.24% more neurons transfected with MAP6 shRNA migrated to bin #5 compared to scramble shRNA transfected neurons. Additionally, 22.33% more scramble shRNA transfected neurons remained in bin #1 compared to MAP6 shRNA transfected neurons. The outcomes indicate that MAP6 depletion accelerates neuronal migration (Figure 3C-E). On the contrary, when tau was depleted via electroporation, neuronal migration was inhibited compared to control mice (Sapir, Frotscher et al. 2012). This indicates that tau and MAP6 have inverse roles in regulating neuronal migration.

**Figure 3:**
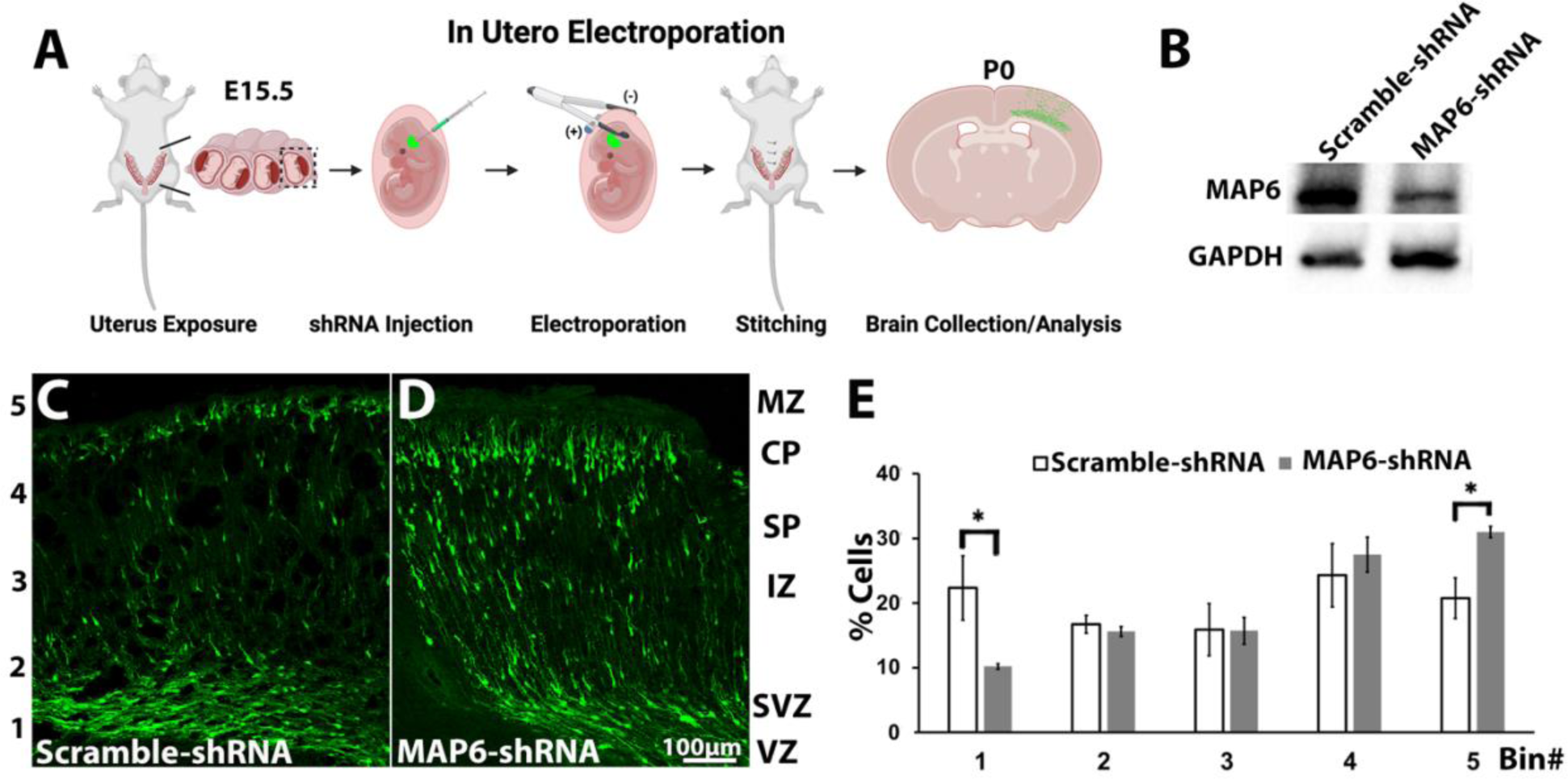
Acceleration of neuronal migration by MAP6 depletion. (A) Schematic of *in utero* electroporation. (B) Representative western blots depicting MAP6 levels in scramble and MAP6 shRNA transfected neurons. (C-D) E15.5 mouse embryos were injected with either scramble shRNA (C) or MAP6 shRNA (D) plasmids, with electroporation facilitating plasmid transfection. The transfected positive neurons are shown in green in the cortex, and the cortex was equally divided into 5 bins. Bin #1 represents the ventricular zone (VZ), and Bin #5 represents the marginal zone (MZ). (E) Graph depicting the percentage of transfected neurons in each bin in the brain of mice transfected with either scramble shRNA or MAP6 shRNA. Bar graph indicates quantification of the percentage of transfected neurons ± SEM in each bin for scramble siRNA (n=5) and MAP6 shRNA (n=5). MAP6 shRNA showed a 12.15% ± 4.12% lower percentage of transfected neurons compared to the scramble (22.33% ± 4.97%) (t-test \**p* < 0.05) in bin #1. MAP6 shRNA showed a 10.24% ± 8.61% higher percentage of transfected neurons compared to the scramble (22.33% ± 4.97%) (t-test \**p* < 0.05) in bin #5. No significant difference (ns) was observed between scramble and MAP6 shRNA in bins # 2, 3, and 4.

**Figure 4:**
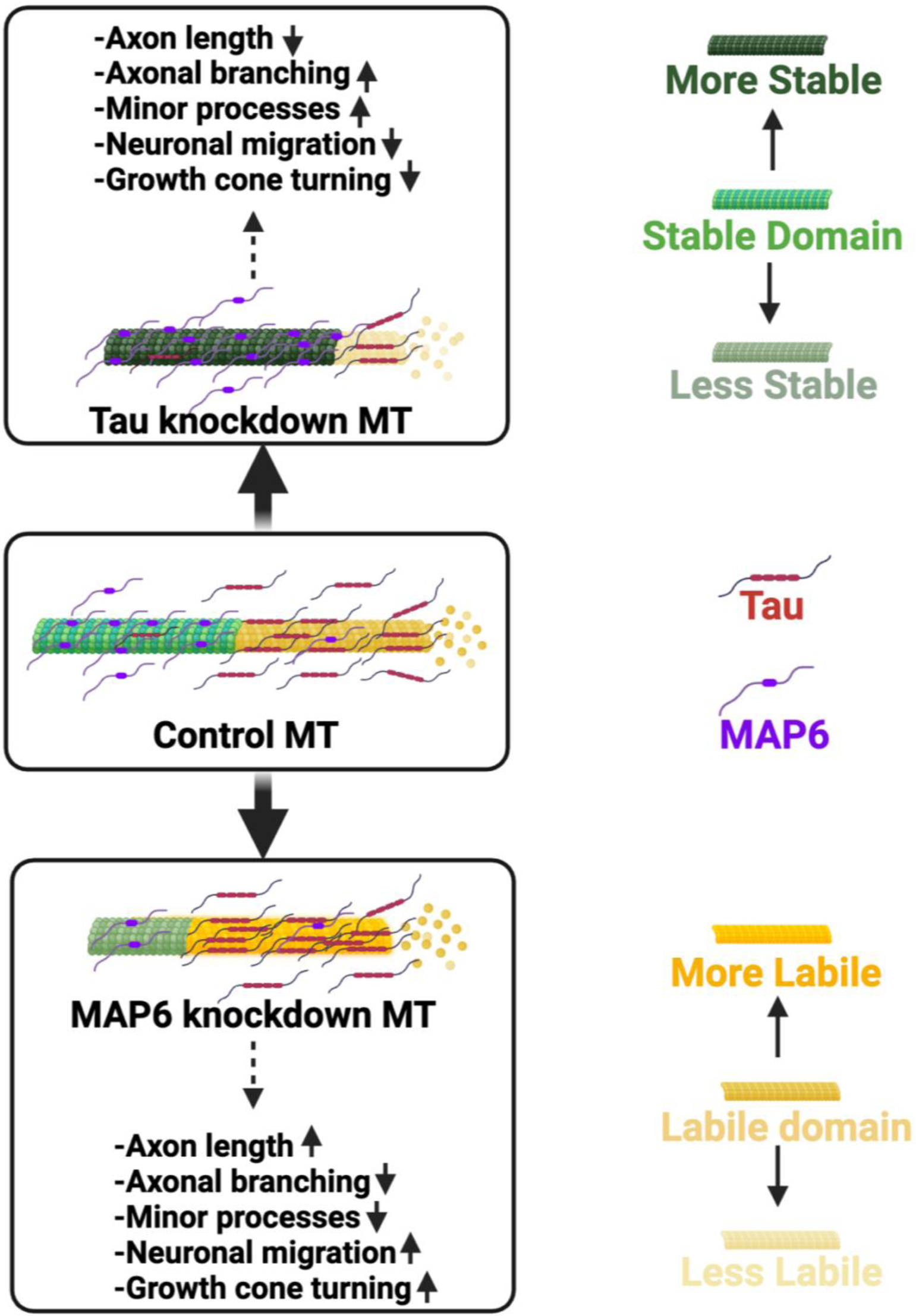
Schematics of how tau and MAP6 are posited to affect aspects of neuronal development and function via their antagonistic impacts on MT dynamics.

### Interpretations and closing remarks

MAP6 and tau, as multifunctional MAPs, orchestrate myriad physiological functions beyond their MT-related activities. While tau has been implicated in synaptic plasticity modulation (Robbins, Clayton et al. 2021), gene expression regulation (Montalbano, Jaworski et al. 2021), signal transduction (Mueller, Combs et al. 2021), and mitochondria homeostasis maintenance (Eckert, Nisbet et al. 2014), MAP6 plays diverse roles in interacting with actin filaments (Peris, Bisbal et al. 2018), membranes (Cuveillier, Delaroche et al. 2020), and neuroreceptors (Cuveillier, Boulan et al. 2021). However, the fact that their knockdown has opposite effects on MT dynamics as well as opposite effects on the various morphological parameters we studied would suggest that those parameters are, in significant part, due to the effects on MT dynamics. This conclusion is fortified by the finding that, with regard to most parameters studied, knocking down both tau and MAP6 together results in preservation of the control phenotypes.

What is the nature of the antagonistic relationship between tau and MAP6? It is unlikely that they compete for the same MT binding site because they have very different MT binding motifs (Cuveillier, Boulan et al. 2021). In fact, some recent studies suggest that MAP6 may bind inside the MT’s lumen in a manner quite distinct from tau (Cuveillier, Delaroche et al. 2020). A potential explanation may lie in a recently proposed mechanism known as MT lattice gating in which the binding of a particular MAP changes the MT lattice to make it more or less amenable to the binding of more of the same MAP or other MAPs (Monroy, Sawyer et al. 2018). Indeed, tau exhibits positive cooperativity in its binding to the MT lattice, wherein the initial tau-MT association induces a conformational change in the lattice. This structural alteration enhances the subsequent binding of tau to the lattice, establishing a positive feedback loop in the tau-MT interaction (Monroy, Sawyer et al. 2018, Siahaan, Tan et al. 2022) which may inhibit MAP6 binds to the MTs. It seems reasonable, based on evidence to date, that tau is usually the better competitor, with MT stabilization occurring only when MAP6 is allowed to outcompete tau, for example through phosphorylation events that decrease tau’s binding affinity to MTs.

Finally, it is worth mentioning that tau has been implicated in a number of neurodegenerative disorders (Arendt, Stieler et al. 2016, Wang and Mandelkow 2016, Gao, Wang et al. 2018, Zhang, Wu et al. 2022, Samudra, Lane-Donovan et al. 2023, Baas, Sullivan et al. 2024) while MAP6 has been implicated in a number of neurodevelopmental disorders (Wei, Sun et al. 2016, Chang, Yang et al. 2018). Our studies suggest that phenotypes resulting from alterations in one of these MAPs may be due in part to corresponding changes in the other MAP, and hence may have implications for both neurodegenerative and neurodevelopmental disorders.

## Materials and methods

### Animals

The use of all animals was in accordance with the guidelines set by NIH and the Drexel University IACUC.

### Rat embryonic hippocampal neuronal cultures

The hippocampus was dissected from TP18-day Sprague Dawley rat fetuses of either sex using cold dissection medium L-15 (Gibco, 21083027). Subsequently, the entire hippocampus was collected and cut into small pieces using microscissors. These pieces were then incubated with 0.25% trypsin and 0.4 mg/ml DNase in a 37-degree water bath for 15 minutes. After incubation, the tissue was washed three times with full plating medium, consisting of neurobasal medium (Gibco, 21103049), 2% B27 supplement (Gibco, A3582801), 0.297% Glucose (Sigma-Aldrich, G8769), 1% GlutaMAX-I (Gibco, 35050061), and 10% fetal bovine serum (Novus Biologicals, S11150-NOV), with each wash lasting 5 minutes. Gentle trituration was performed to separate any clumps, and the neurons were then counted using a hemocytometer. The neurons were cultured for 2 days and then replated for an additional 2 days of culture.

### Rat superior cervical ganglion (SCG) neuronal cultures

SCGs obtained from postnatal day 0 (P0) to P3 Sprague Dawley rat pups of both sexes and cultured with established protocols (Muralidharan and Baas 2019). Prior to plating, targeted siRNA, as described below, was introduced into SCG neurons through nucleofection using the Nucleofector 2b system from Lonza. After nucleofection, the cells were cultured for a period of 48 hours on 35-mm-diameter culture dishes coated with 0.1 mg/ml poly-D-lysine (Sigma Aldrich, P0899-50MG). Subsequently, the cells were transferred to glass-bottomed dishes (Cellvis, D35-14-1.5-N), which were also coated with 0.1 mg/ml poly-D-lysine, and a 30 µl droplet of laminin (Invitrogen, 23017-015) was applied. 50,000 SCG neurons were replated on to each dish, and strategically placed in the region where the laminin droplet had been deposited. On day 4, these neurons were fixed and subjected to immunostaining with ßIII-tubulin, Laminin, and DAPI, as described below.

### RNAi – based knockdown of tau and MAP6

Small interfering RNA (siRNA) targeting scramble, tau, or MAP6 was introduced into rat hippocampal or SCG neurons using a Nucleofector (Amaxa), and the manufacturer’s G-13 program. The tau or MAP6 siRNA consisted of a mixture of three distinct siRNA duplexes designed to target various regions of each molecule, acquired from Sigma’s custom siRNA service. These are the same sequences verified in our previous study in which we conducted appropriate controls, including rescue experiments (Qiang, Sun et al. 2018). A nonspecific duplex III, previously utilized, served as the control. The siRNA was prepared at a concentration of 200 nM, aliquoted, and subsequently stored at −20°C. The final siRNA concentration utilized for transfection was 10 nM. Subsequent to the transfection, the neurons were cultured for 48 hours on plastic dishes coated with either poly-L-lysine (Sigma Aldrich, P2636-25MG) for hippocampal neurons or poly-D-lysine for SCG neurons before undergoing replating. The siRNA-treated neurons were then maintained for 4 days prior to fixation.

### MAP6 shRNA design and validation

For making pCSV2-Venus-mouse MAP6 shRNA, the sequences below were synthesized, for the MAP6_shRNA_forward: gatcc (BamHI)/GACCTCAACGAGCCATAAATTCAAGAGATTTATGGCTCGTTGAGGTCTTTTT GGAA/a. For the MAP6_shRNA_reverse: agctt (Hind III)/TTCCAAAAAGACCTCAACGAGCCATAAATCTCTTGAATTTATGGCTCGTTGAGGTC/g. The annealed oligonucleotides were ligated into the BamHI/HindIII-digested pSCV2-Venus vector. The MAP6 shRNA target 1545-1563 (19 mer). Then the MAP6 shRNA efficiency was validated in hippocampal neurons. Briefly, either 3.6 µg of scramble shRNA or MAP6 shRNA were transfected into hippocampal neurons using Lipofectamine 2000 transfection reagent (Invitrogen, 11668030). Neurons were then collected for protein analysis using western blot.

### Western Blotting (WB)

The cell lysates were collected based on the protocol mentioned in the previous study (Qiang, Sun et al. 2018). In short, WB was performed using the Bio-Rad system. In essence, 20 µg of protein was loaded into each of the 10 wells on the WB gel, and the gel was run under 100 volts for 2 hours. Subsequently, the gel was transferred at 30 volts overnight. Following this, the membranes were incubated with primary antibodies specific to tau (R1) (from. Dr. Kanaan, 1:2000), MAP6 (from Dr. Andrieux, 1:600), tyrosinated-tubulin (Sigma-Aldrich, MAB1864, 1:1000), or GAPDH (Abcam, ab8245, 1:2000), followed by HRP secondary antibodies (Qiang, Sun et al. 2018).

### Immunocytochemistry (ICC)

Both hippocampal and SCG neurons were fixed using a co-fixative buffer comprising 4% paraformaldehyde (Sigma Aldrich, 158127-3KG), 1X PHEM buffer (containing PIPES, HEPES, EDTA, MgCl2), 0.2% glutaraldehyde (Electron Microscope Science, 16019), and 0.1% Triton X-100 (Sigma Aldrich, 122H0766) for 10 minutes at room temperature, followed by three washes with PBS. Subsequently, immunochemistry was conducted following previously established protocols (Baas and Qiang 2019).

### *In Utero* Electroporation (IUE)

This methodology was executed following established protocols (Bennison, Blazejewski et al. 2023). Briefly, pregnant E15.5 mice were anesthetized, and the uterine horn was exposed. Utilizing a pulled-glass micropipette, 1-2 µl of either scramble or MAP6 shRNA plasmids were injected into the lateral ventricle. Following this, electric pulses (consisting of three pulses at 32 V each) were administered via tweezers-type electrodes placed over the uterine muscle, using the CUY21SC electroporator from Nepa GENE. Subsequently, the uterine horn was carefully repositioned within the abdomen, allowing uninterrupted embryonic development. Venus fluorescent protein encoded in pSCV2-Venus vector was used to visualize electroporated neurons.

### Immunohistochemistry (IHC)

The brains of P0 mouse were dissected and fixed in a 4% paraformaldehyde for 24 hours at 4 degrees. Subsequently, the brains were transferred to a 25% sucrose solution to undergo dehydration and facilitate sinking. Cryosectioning was carried out to obtain brain slices with a thickness of 25 µm, with the brains embedded in the M1 matrix (Epredia, 1310). These brain slices were allowed to air-dry overnight and then stained with 4’,6-Diamidino-2-Phenylindole, Dihydrochloride (DAPI) (Invitrogen, D1306, 1:20,000) for 5 minutes. Finally, the stained brain slices were mounted using fluoro-gel mounting medium (Electron Microscopy Sciences, 1798510) and were ready for imaging.

### Imaging, statistical analysis and data processes

Imaging was conducted utilizing either the Leica True Confocal System SP8 for navigation and Z-stack acquisition or the Zeiss AxioObserver. Quantitative analysis was carried out using either Zeiss blue edition software or Fiji software. The acquired data was recorded in Excel, and subsequent bar graphs were generated using GraphPad. For statistical analysis, a t-test was employed to compare the average mean values between two comparison groups, while one-way ANOVA was used when comparing the average mean values across more than two comparison groups. The data are presented as Mean ± SEM, statistical significance is denoted on figures in the following manner: \**p* < 0.05, ‘ns’ represents no significant difference. The figures and graphs were organized using Adobe Photoshop 2023.

## Acknowledgments

We thank Dr. Nicholas M. Kanaan for Tau (R1) antibody, and Dr. Annie Andrieux for MAP6 antibody and Shirin Kaye for quantifying the neuronal migration data.

